# The impact of behavioral interventions on co-infection dynamics: an exploration of the effects of home isolation

**DOI:** 10.1101/358093

**Authors:** Diana M Hendrickx, Steven Abrams, Niel Hens

**Affiliations:** Center for Statistics, Interuniversity Institute for Biostatistics and statistical Bioinformatics, Hasselt University, Diepenbeek, Belgium; Centre for Health Economics Research and Modelling Infectious Diseases, Vaccine and Infectious Disease Institute, University of Antwerp, Antwerp, Belgium

## Abstract

Behavioral changes due to the development of symptoms have been studied in mono-infections. However, in reality, multiple infections are circulating within the same time period and behavioral changes resulting from contraction of one of the diseases affect the dynamics of the other.

The present study aims at assessing the effect of home isolation on the joint dynamics of two infectious diseases, including co-infection, assuming that the two diseases do not confer cross-immunity. We use an age- and time- structured co-infection model based on partial differential equations. Social contact matrices, describing different mixing patterns of symptomatic and asymptomatic individuals are incorporated into the calculation of the age- and time-specific marginal forces of infection.

Two scenarios are simulated, assuming that one of the diseases has more severe symptoms than the other. In the first scenario, people stay only at home when having symptoms of the most severe disease. In the second scenario, twice as many people stay at home when having symptoms of the most severe disease than when having symptoms of the other disease.

The results show that the impact of home isolation on the joint dynamics of two infectious diseases depends on the epidemiological parameters and properties of the diseases (e.g., basic reproduction number, symptom severity). In case both diseases have a low to moderate basic reproduction number, and there is no home isolation for the less severe disease, the final size of the less severe disease increases with the proportion of symptomatic cases of the most severe disease staying at home, after an initial decrease. When twice as many people stay at home when having symptoms of the most severe disease than when having symptoms of the other disease, increasing the proportion staying at home always reduces the final size of both diseases, and the number of co-infections.

In conclusion, when providing advise if people should stay at home in the context of two or more co-circulating diseases, one has to take into account epidemiological parameters and symptom severity.

## 1 Introduction

Jointly modeling the dynamics of two or more infectious diseases with or without similar transmission routes can provide new insights in the interaction among these different pathogens [5, 7, 8, 12]. For airborne diseases, deterministic compartmental models described by ordinary differential equations (ODEs) have been proven to provide a suitable mathematical framework for studying such interactions [7, 8]. Such ODE-based co-infection models typically describe the transmission dynamics of two (or more) infectious diseases, and the flow of individuals between different compartments or states (e.g., susceptible, infected, recovered), in function of calendar time. Alternatively, age-specific effects could be studied, at least when assuming endemic equilibrium for the infections at hand similarly [6, 9].

Apart from calendar time, age is also an important factor influencing the dynamics of infectious diseases. Within the same calendar year, transmission parameters can differ for people of various ages, e.g., for childhood diseases the infection risk tends to be lower for adults and elderly as compared to children. Hence, compartmental models including both calendar time and age effects provide a straightforward extension of the aforementioned models. The flow of individuals in such models is then described using a system of partial differential equations (PDEs) in time and age. Age structure can be included in the model via contact or mixing matrices, including social contact rates among individuals in different age categories, the population age distribution and age-specific mortality rates [2, 6].

In addition to age and calendar time effects, implicitly decomposing the population in various subgroups, one can decompose the subpopulation of infectious individuals further into symptomatic and asymptomatic cases, which makes sense if for the pathogens under study the occurrence of asymptomatic infections is agreed upon. While most individuals change their social contact behavior when experiencing symptoms [3], at least when these symptoms are moderate to severe, by staying at home, asymptomatic individuals will show similar contact patterns as compared to individuals who are uninfected (either susceptible or immunized). Furthermore, symptomatic individuals are presumed to be more contagious than asymptomatic individuals, which has been demonstrated in the context of influenza by Van Kerckhove et al. [14]. Behavioral changes due to the development of symptoms and differences in contagiousness between symptomatic and asymptomatic people have recently been implemented and the effects thereof have been studied using compartmental models for mono-infections [10]. More specifically, these authors [10] showed that in case of influenza-like illness (ILI), the total number of cases can be reduced by 39% or 63% when 50% or all symptomatic individuals, respectively, would stay at home immediately after the onset of symptoms.

The present study extends the work by Santermans et al. [10] in the sense that our approach incorporates social contact matrices for both symptomatic and asymptomatic individuals, together with differences in infectiousness among those two groups, in an age- and time-structured co-infection model for two diseases which is described using a system of PDEs. We assume that there is no cross-immunity induced for the diseases at hand. First, we have studied the effect of staying at home when having symptoms for one disease on the final size of the other infection. More specifically, we studied how the following infectious disease parameters influence this effect: basic reproduction numbers, infectious period, fraction of symptomatic cases, number of contacts and the delay between the two epidemic outbreaks. Second, we studied two diseases with different symptom severity, where twice as many people stayed at home when having symptoms of the most severe disease than when having symptoms of the other disease. Both the basic reproduction number and the proportion staying at home were varied.

The paper is organized as follows. In Section 2, we describe the co-infection model configuration, parameter settings, and the scenarios and model variations considered. In Section 3, the results from investigating the effect of behavioral changes due to having symptoms on the model output are presented. Section 4 discusses our main findings and summarizes our conclusions and recommendations for further research.

## 2 Methods

### 2.1 Co-infection model setup

The co-infection model used in this paper is an age-structured Susceptible-Infected-Recovered (SIR) compartmental transmission model, describing the joint disease dynamics with regard to two immunizing infections conferring lifelong humoral immunity. The model was implemented in R3.1.1 and R3.3.2 using the deSolve package [13]. In total, the co-infection model consists of 9 different compartments or states, which are described in detail in Table 1. Figure 1 shows a schematic diagram depicting the different compartments and the flow of individuals between the states in the model. In particular, these flows can be described using a system of PDEs, which is available in Appendix A.

**Table 1.** Compartments in the SIR model used in this study.

| State | Meaning |
| --- | --- |
| $S_{12}$ | susceptible for both infections |
| $I_{1S}$ | infected by pathogen 1, susceptible for infection by pathogen 2<br>$I_{1Sa}$: asymptomatic infection; $I_{1Ss}$: symptomatic infection |
| $I_{S2}$ | infected by pathogen 2, susceptible for infection by pathogen 1<br>$I_{S2a}$: asymptomatic infection; $I_{S2s}$: symptomatic infection |
| $I_{12}$ | co-infection<br>$I_{12aa}$: asymptomatic; $I_{12as}$: only symptoms of disease 2;<br>$I_{12sa}$: only symptoms of disease 1; $I_{12ss}$: symptoms of both diseases |
| $I_{1R}$ | infected by pathogen 1, recovered from infection by pathogen 2<br>$I_{1Ra}$: asymptomatic infection; $I_{1Rs}$: symptomatic infection |
| $I_{R2}$ | infected by pathogen 2, recovered from infection by pathogen 1<br>$I_{R2a}$: asymptomatic infection; $I_{R2s}$: symptomatic infection |
| $R_{1S}$ | recovered from infection by pathogen 1, susceptible for infection by pathogen 2 |
| $R_{S2}$ | recovered from infection by pathogen 2, susceptible for infection by pathogen 1 |
| $R_{12}$ | recovered from both infections |

**Figure 1.**
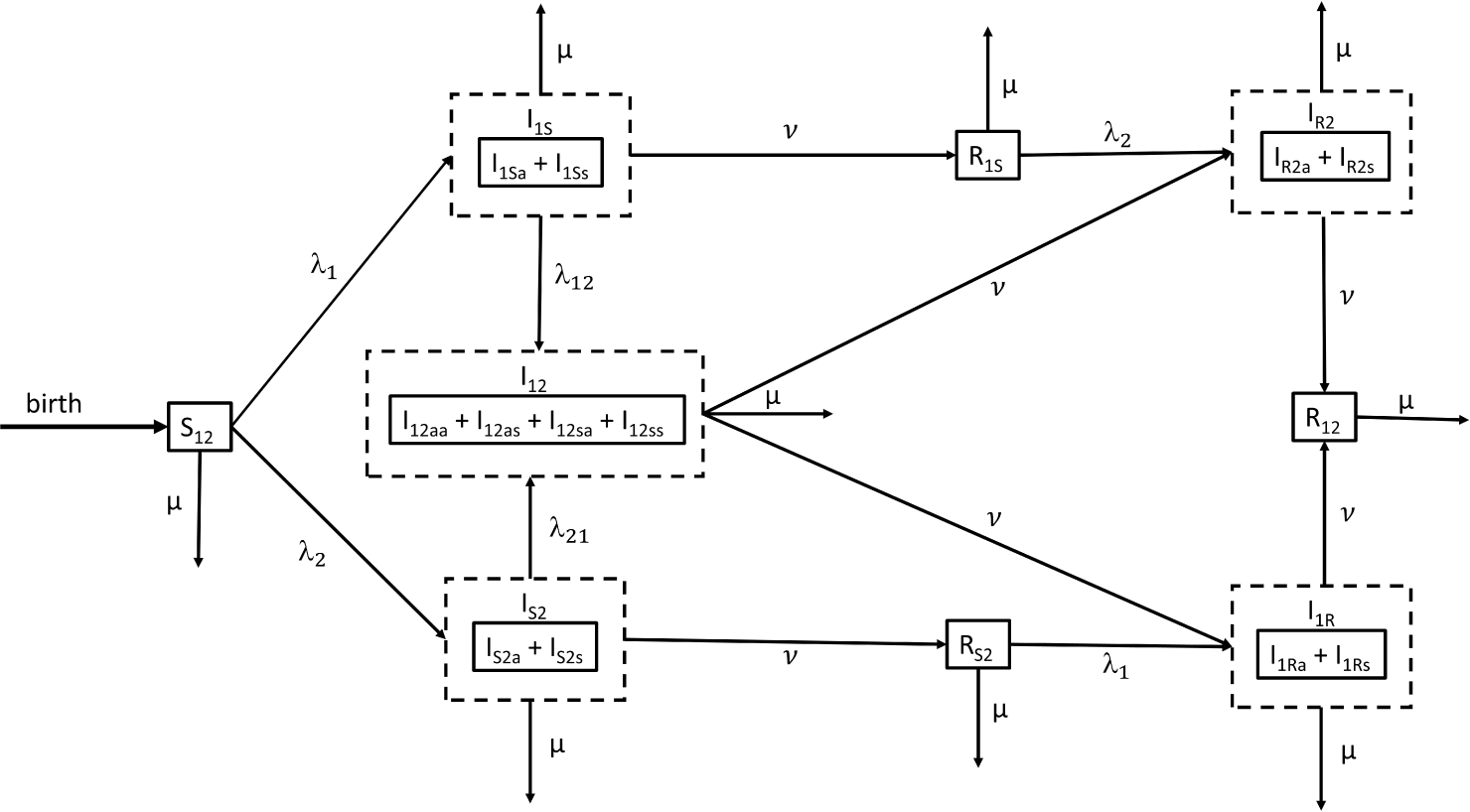
Schematic diagram of the SIR model used in this paper. Model parameters are *µ*: natural mortality rate; λ_1_: the marginal force of infection for infection 1;λ_2_: the marginal force of infection for infection 2; λ_12_: the force of infection for infection 2, conditional on infection 1; λ_21_: the force of infection for infection 1, conditional on infection 2; *ν*: the recovery rate (similar for infection 1 and 2)

### 2.2 Parameter configuration

#### 2.2.1 Age structure

In the simulations, individuals in the population with ages ranging from 0 to 85 years in the different compartments are divided into one year age categories. Model parameters are assumed to be age-specific and are allowed to differ by calendar time (see also next paragraph). However, to summarize the results in this manuscript, individuals are grouped into two age categories: individuals of 0 – 18 years and 19 – 85 years.

#### 2.2.2 Social contact matrices

The age- and time-specific marginal and conditional forces of infection (FOI)λ_1_(*a,t*), λ_2_(*a,t*), λ_12_(*a,t*), and λ_21_(*a,t*), were related to the social contact data using the mass action approach by Wallinga and colleagues [15] (see, e.g., A for more details). The corresponding values of the FOI were calculated for various hypothesized values of the basic reproduction number *R*_0_, that is, the average number of secondary infections produced by a single ‘typical’ infectious individual during his/her entire infectious period when introduced in a fully susceptible population. In this study, four different mixing matrices were constructed based on social contact survey data:

- *C_aa_*: asymptomatic mixing matrix describing the age-specific mixing behavior of asymptomatic individuals;
- *C_sa_* and *C_as_*: mixing matrix for individuals only having symptoms of disease 1 resp. disease 2;
- *C_ss_*: mixing matrix when having symptoms of both diseases under study.

Data from the social contact survey studied in Van Kerckhove et al. [14] were used to construct 2*×* 2 contact matrices *C^A^*, *C^S^* and 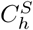 for the two age categories defined previously. *C^A^* is the asymptomatic contact matrix, which is assumed to be the same as the contact matrix for ‘healthy’ individuals (i.e., ‘healthy’ with regard to the infections at hand). Furthermore, *C^S^* is the contact matrix for symptomatic individuals not staying at home and 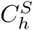 the contact matrix for symptomatic individuals staying at home.

Let *p*_1_ and *p*_2_ represent the proportions of individuals staying at home when having symptoms of disease 1 and 2, respectively, and let *p*_12_ be the proportion of individuals staying at home when having symptoms of both diseases. In the first scenario, where people only stay at home for the most severe disease, we assume that *p*_12_ = *p*_1_. In the second scenario, where people stay at home for both diseases, we assume that *p*_12_ will be larger than *p*_1_ and *p*_2_. In this study, we define *p*_12_ as *p*_1_ + *p*_2_ *− p*_1_*p*_2_, so that *p*_12_ is always the largest of the three proportions *p*_1_, *p*_2_ and *p*_12_. According to the aforementioned notation, the social contact matrices are given by: *C_aa_ = C^A^*; 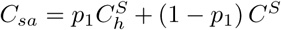; 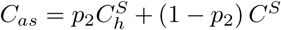; and 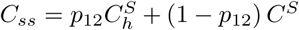.

#### 2.2.3 Model variations and scenarios

In a first scenario, simulations were run for two infections starting at the same time with an infectious period of 7 days and 60% of infections being symptomatic. Symptomatic cases were supposed to be three times as infectious as asymptomatic cases. The basic reproduction number *R*_0_ was varied between 1.5 and 6.5, with steps of size 1. The percentage of individuals staying at home when having symptoms of disease 1 was varied between 0% and 100%, with steps of size 5%. People were supposed not to stay at home when having symptoms of the second disease. The following model variations were applied to this scenario:

- infectious period of respectively 14 days and 21 days;
- symptomatic cases are six times (respectively nine times) as infectious as asymptomatic cases;
- 90% (respectively 30%) of the infected individuals are symptomatic;
- symptomatic individuals not staying at home and asymptomatic individuals would have the same mixing patterns as uninfected individuals;
- a difference of 0.3 between the basic reproduction numbers of the two diseases;
- a delay of one month between the two diseases.

As a second scenario, two infections were studied, for which the proportion staying at home when having symptoms of the less severe disease was half of the proportion staying at home when having symptoms for the other disease.

For all scenarios in which both infections are introduced simultaneously, the model was initialized with 1 co-infected person of 10 years old and the remainder of the population was considered susceptible for both infections. For the scenarios with a delay between the starting times of the two infections, the start of infection 1 (resp. infection 2) was initialized with 1 person of 10 years old, mono-infected by pathogen 1 (resp. pathogen 2) and still susceptible for the other infection.

### 2.3 Solving the system of PDEs – Method of lines

In order to numerically solve the system of PDEs for the co-infection model presented in Figure 1, we rely on the method of lines [11] in which the age dimension is discretized and only the time dimension remains continuous. Consequently, the method of lines leads to a system of ODEs that can be solved by means of a numerical method for initial value ODEs. For more details regarding the method of lines, we refer to [11].

## 3 Results

In this section, we discuss the results of our simulation approach. First, we investigated the effect of staying at home when having symptoms of one disease on the dynamics of the other infection. Second, the influence of the following model parameters on the observed effect was studied: the basic reproduction number, the infectious period, the infectiousness of symptomatic versus asymptomatic individuals, the proportion of cases being symptomatic, the number of contacts and delays between the start of the two infections. Third, the effect of staying at home for two diseases where twice as many people stay at home when having symptoms of the most severe disease compared to the other disease was investigated.

The observed results are explained by comparing the dynamic profiles of the infections, including the peak time of infection.

### 3.1 Influence of home isolation when having symptoms of one disease

Changes in contact behavior by staying at home when having symptoms of the most severe disease (disease 1) induces the final size of co-infection to decrease (Figure A.1). Figure A.2 graphically depicts the effect of *R*_0_ on the total number of co-infections for a range of *R*_0_ values between 1 and 1.5. Here, we observe that staying at home counteracts the natural increase of the final size of co-infections with increasing *R*_0_.

For infection 2, different scenarios can be observed, depending on the value of the basic reproduction number. Figure 2 depicts the final size of infection 2 for varying percentages of individuals staying at home when having symptoms of disease 1 (*p*_1_ ranges from 0% up to 100% in steps of size 5%), and varying values of the basic reproduction number (*R*_0_ ranges from 1.5 up to 6.5 in steps of size one). For small to moderate values of *R*_0_ the final size of infection 2 tends to increase, after an initial decrease for *R*_0_ = 1.5, 2.5, 3.5, 4.5 and 5.5, with an increasing value of *p*_1_ (for *R*_0_ = 1.5, see also Figure A.3). The value of *p*_1_ corresponding with the minimal final size of infection 2 increases with increasing *R*_0_. However, for high *R*_0_ values, hence, more contagious pathogens, the final size of infection 2 decreases with increasing *p*_1_ values.

**Figure 2.**
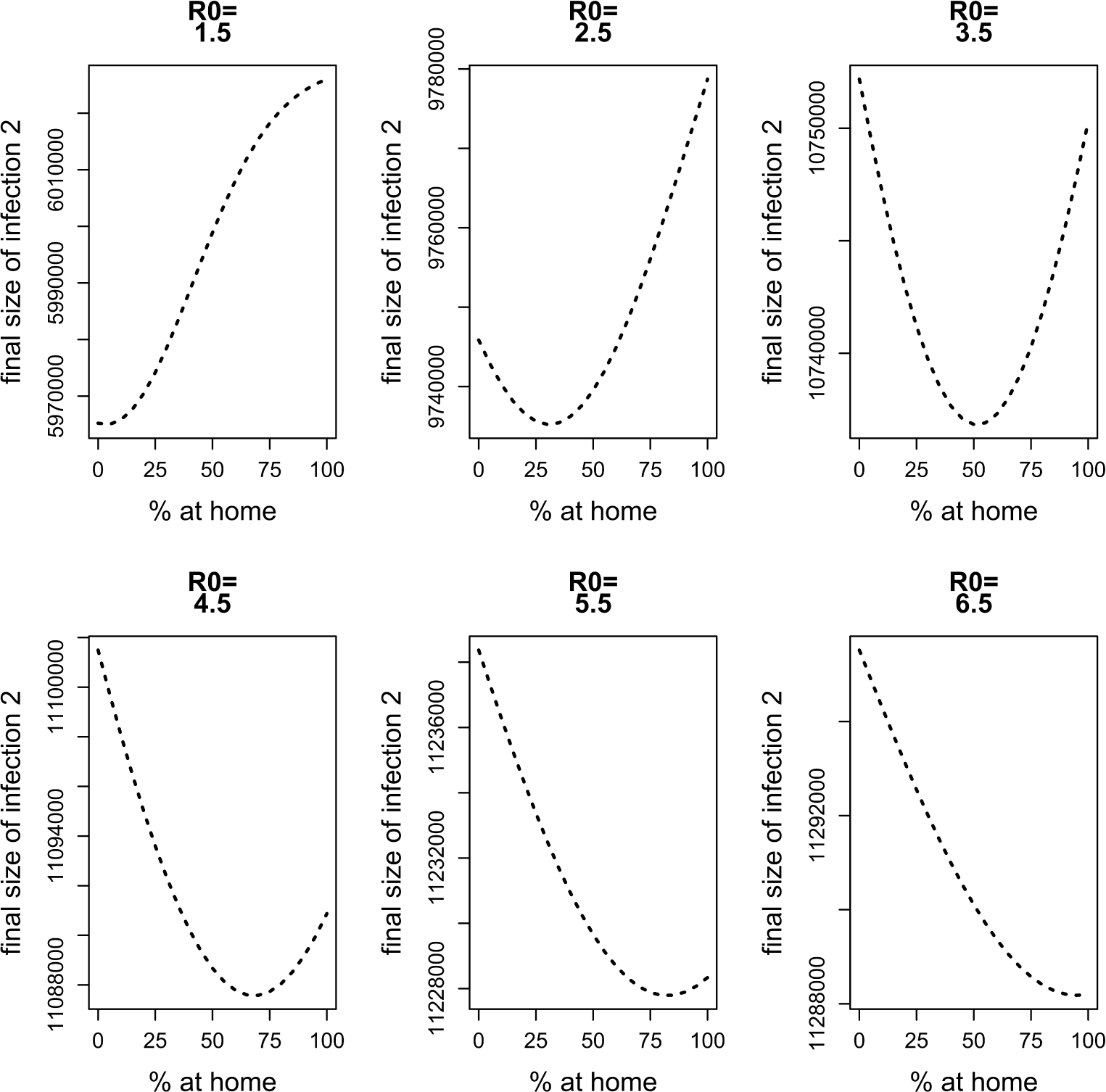
Final size of infection 2 against the percentage staying at home when having symptoms of disease 1 for different values of *R*_0_.

From Figure 3 (upper panel), it can be observed that staying at home when having symptoms of the most severe infection (infection 1) leads to a shift of the peak time of new infections with pathogen 1. This shift of the peak time increases with increasing *p*_1_ values. As a consequence, at the peak time of the less severe disease (infection 2), which equals the peak time of infection 1 in case of no home isolation (solid line Figure 3, upper panel), the number of symptomatic cases of infection 1 decreases with increasing *p*_1_ (middle panel). Furthermore, the number of people staying at home at the peak time of infection 2 decreases with increasing *p*_1_ (lower panel). This means that the two groups of individuals that have less contacts (individuals staying at home and symptomatic cases not staying at home) represent a smaller fraction of the population. As a consequence, people will have on average more contacts and will have a higher probability to acquire infection 2. This explains the increasing trend in the final size of infection 2.

**Figure 3.**
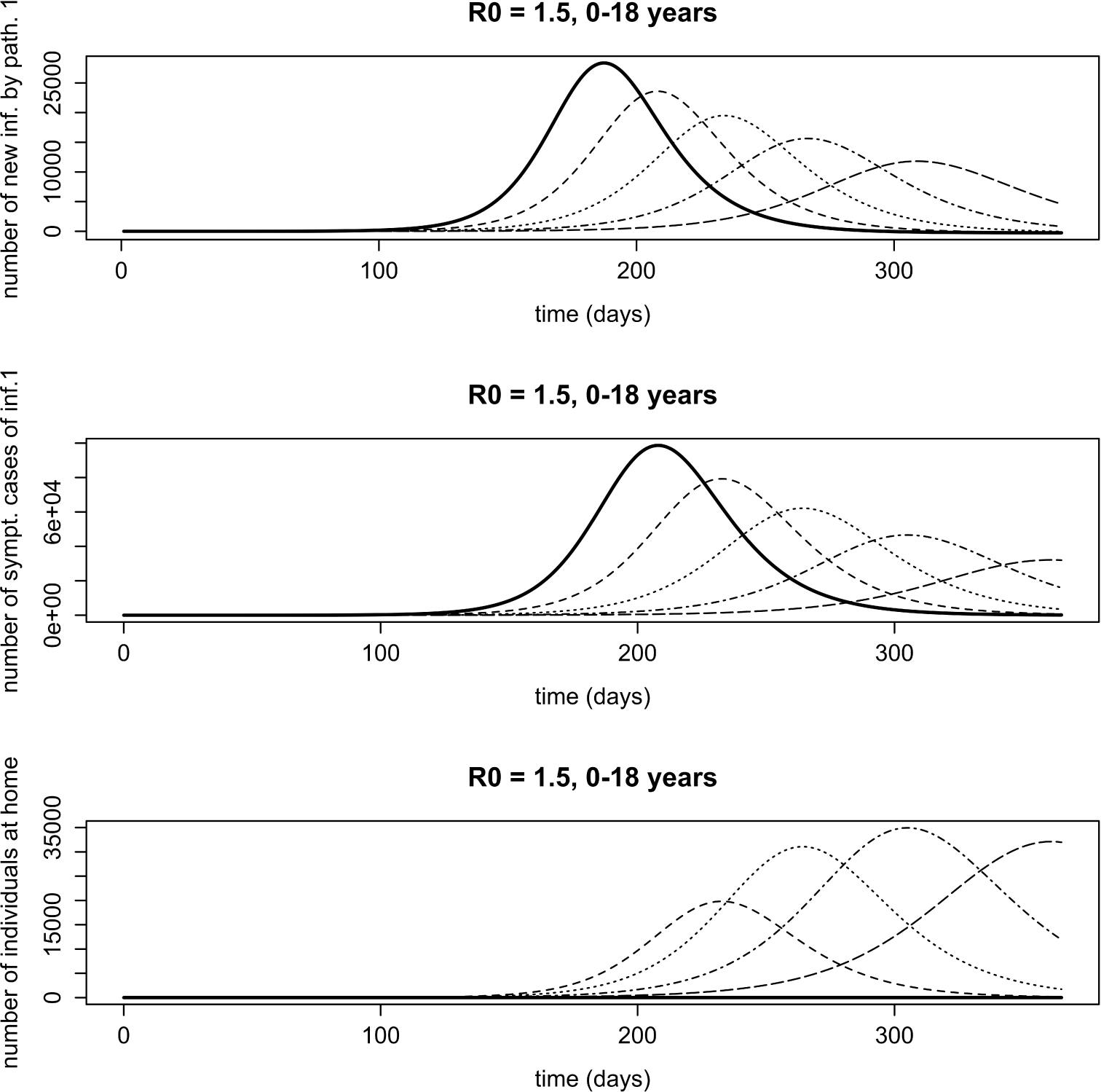
Influence of home isolation for disease 1 only, for age class 0 − 18 years in case of two infections with an infectious period of 7 days and *R*_0_ = 1.5. Solid line: 0% at home, dashed line: 25% at home, dotted line: 50% at home, dotdashed line: 75% at home, longdashed line: 100% at home. Upper figure: number of new infections with pathogen 1. Because there is no home isolation for disease 2, the number of new infections with pathogen 2 coincides with the solid line; middle figure: number of symptomatic cases of infection 1; lower figure: number of symptomatic cases of infection 1 staying at home.

When comparing Figure 3 with Figure A.4, it can be observed that similar scenarios occur for both age classes.

For larger values of *R*_0_ (e.g. *R*_0_ = 3.5), the shift of the peak time of new infections with pathogen 1 becomes smaller (see Figure A.5). At the peak time of infection 2, the decrease of the number of people staying at home occurs over a smaller interval (see Figure A.5). As a consequence, the increasing part of the graph of the final size of infection 2 in Figure 2 becomes smaller with increasing *R*_0_ values.

### 3.2 Influence of model parameters on the observed effects

Figures A.7–A.9 show that the following model parameters have little or no influence on the effects observed in Section 3.1: the infectious period, the infectiousness of symptomatic versus asymptomatic cases, the fraction of symptomatic cases.

If symptomatic cases of infection 1 not staying at home and asymptomatic cases would have the same mixing patterns, the final size of infection 2 will never be higher when staying at home than without home isolation. The interval that shows an increasing final size of infection 2 with increased home isolation becomes smaller (see Figure A.10).

When *R*_0_ of pathogen 1 is smaller (resp. larger) than *R*_0_ of pathogen 2, a decrease of the final size of infection 2 with *p*_1_ starts to occur at higher (resp. lower) values of *R*_0_ for pathogen 2, compared to two pathogens with equal *R*_0_ (Figure A.11).

In case infection 1 starts earlier (resp. later) than infection 2, a decrease of the final size of infection 2 with *p*_1_ starts to occur at smaller (resp. larger) *R*_0_ (Figure A.12), compared to two infections with equal starting time.

### 3.3 Influence of symptom severity

A more realistic setting, in which twice as many people stay at home when having symptoms of the most severe disease (disease 1) than when having symptoms of the less severe disease (disease 2)(*p*_1_ = 2 · *p*_2_), was simulated. More specifically, the following scenario was studied in detail:

- Based on the 2009 A/H1N1pdm influenza epidemic, we assume that he percentage staying at home when having symptoms of the most severe disease (*p*_1_) is 70% (Kim Van Kerckhove, personal communication).
- The percentage staying at home when having symptoms of the less severe disease (*p*_2_) is 35%.
- Symptomatic individuals are three times as infectious as asymptomatic individuals [14].
- For both diseases, 66% of the infections are symptomatic [14].
- Both diseases have *R*_0_ = 1.5, an infectious period of 7 days and start at the same time.

Figure 4 shows that for infection 1, the numbers of susceptible and infected are almost equal when not staying at home compared to the situation in which 35% of symptomatic individuals having disease 2 stay at home (and there is no home isolation for disease 1). A higher number of susceptible and a lower number of infected individuals are observed when 70% of symptomatic individuals with disease 1 stay at home (and there is no home isolation for disease 2). Furthermore, the peak time of infection shifts to the right (the peak of epidemic 1 is delayed). A similar scenario is observed when 70% of symptomatic individuals with disease 1 and 35% of symptomatic individuals with disease 2 stay at home.

**Figure 4.**
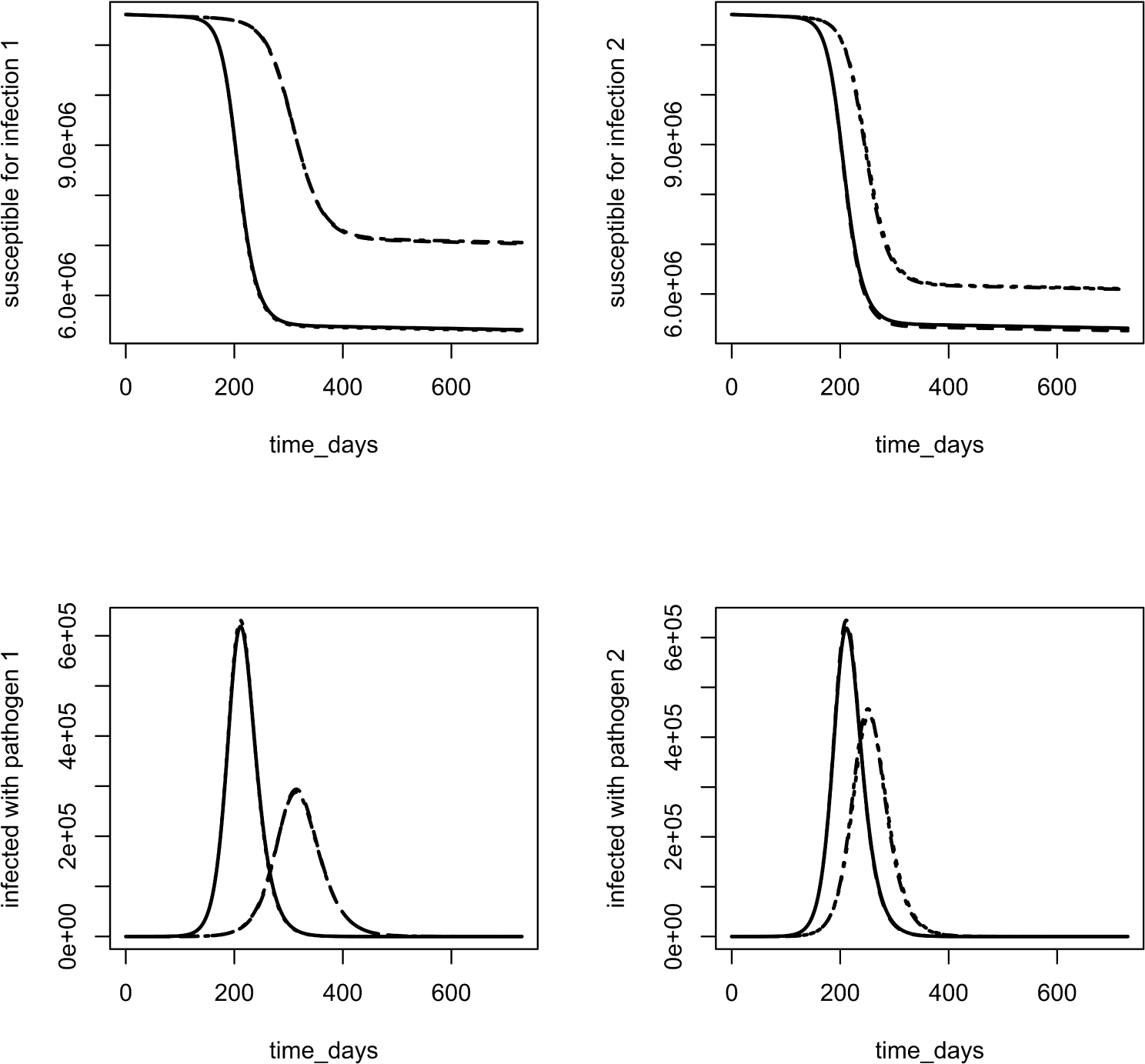
First row: number of susceptible for disease 1 (left) and disease 2 (right); second row: number of infected with disease 1 (left) and disease 2 (right). Scenarios: no home isolation (solid); 70% symptomatic cases at home for disease 1, no home isolation for disease 2 (dashed); 35% symptomatic cases at home for disease 2, no home isolation for disease 1 (dotted); 70% symptomatic cases at home for disease 1, 35% symptomatic cases at home for disease 2 (dotdashed). In the left figures, the following lines coincide: solid and dotted; dashed and dotdashed. In the right figures, the following lines coincide: solid and dashed; dotted and dotdashed.

For disease 2, Figure 4 shows that the number of susceptible (resp. infected) is a bit lower (resp. higher) when 70% symptomatic individuals with disease 1 stay at home (and there is no home isolation for disease 2) than when there is no home isolation for both diseases. A significantly higher (resp. lower) number of susceptible (resp. infected) is observed when 35% of symptomatic individuals with disease 2 stay at home (and there is no home isolation for disease 1). Furthermore, the peak time of infection shifts to the right (the peak of epidemic 2 is delayed). When 70% of symptomatic individuals with disease 1 and 35% of symptomatic individuals with disease 2 stay at home, the number of susceptible (resp. infected) is a bit lower (resp. higher) than when only 35% of symptomatic individuals with disease 2 stay at home (and there is no home isolation for disease 1).

Figure 5 shows the number of individuals recovered from infection 1, infection 2 and co-infections. Like mentioned before, staying at home when having symptoms of only one disease has a significant positive effect on that disease, and a slightly negative effect on the other. When considering co-infections, the most advantageous scenario is staying at home when having symptoms of the most severe disease, followed by staying at home when having symptoms of one of the two diseases.

**Figure 5.**
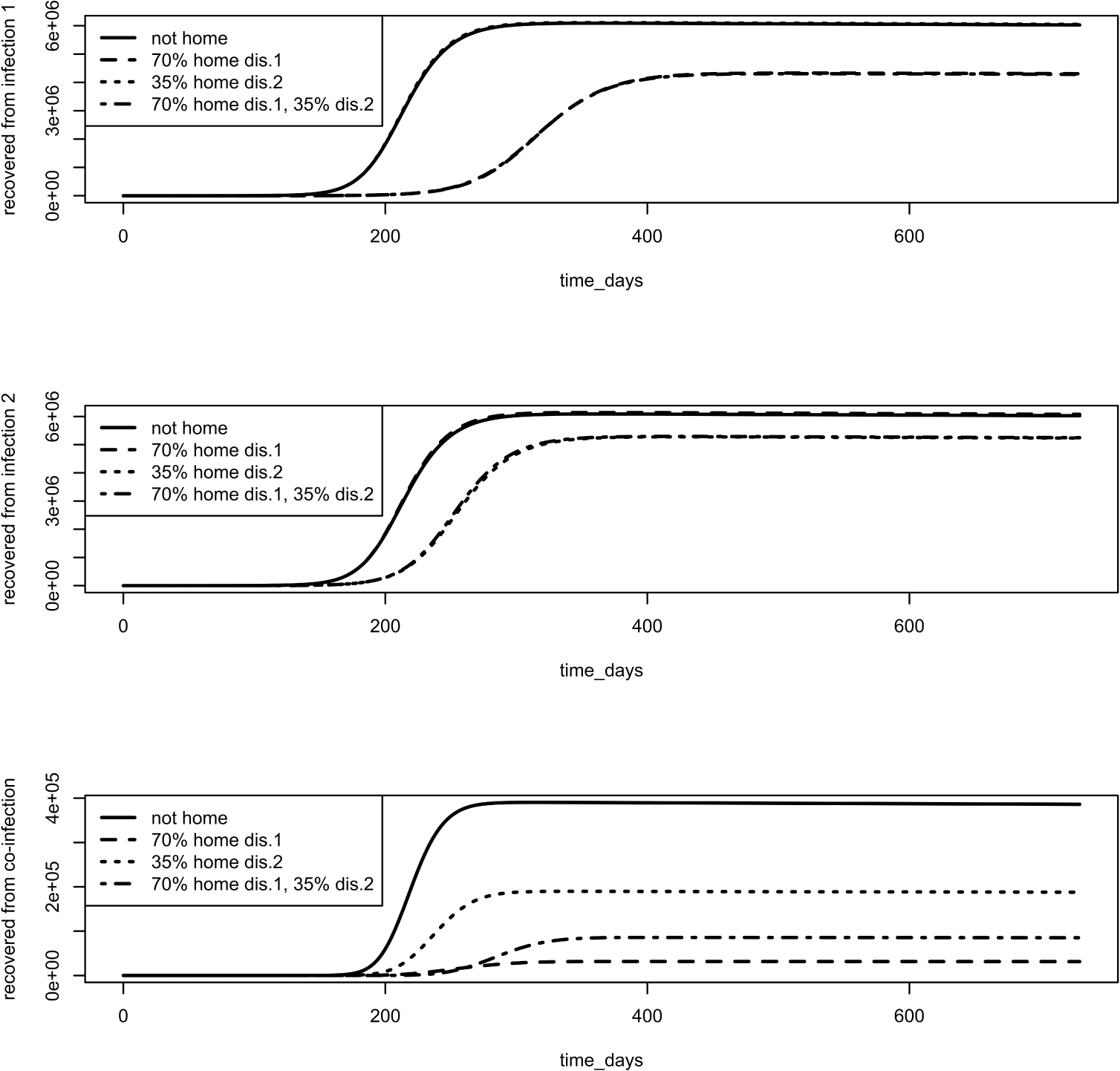
Number of people recovered from disease 1 (upper panel), disease 2 (middle panel) and co-infection (lower panel). Scenarios: no home isolation (solid); 70% symptomatic cases at home for disease 1, no home isolation for disease 2 (dashed); 35% symptomatic cases at home for disease 2, no home isolation for disease 1 (dotted); 70% symptomatic cases at home for disease 1, 35% symptomatic cases at home for disease 2 (dotdashed). In the upper panel, the following lines coincide: solid and dotted; dashed and dotdashed. In the middle panel, the following lines coincide: solid and dashed; dotted and dotdashed.

Figure A.13 shows that in case twice as many people stay at home when having symptoms of the most severe disease than when having symptoms of the other, increasing *p*_1_ and *p*_2_ always decreases the final size of disease 1, disease 2 and co-infections, irrespective of the value of *R*_0_.

## 4 Discussion

In this paper, we explored various scenarios of altering behavior, upon contraction of an infection, using a co-infection model. More specifically, we studied the effect of changing social contact behavior on the dynamics and final size of emerging infections, aiming at an improved understanding of social interventions distancing. When studying an influenza-like disease in isolation, Santermans et al. [10] showed that staying at home leads to a significant reduction of the final size of the disease. However, multiple infectious diseases often circulate within the same period, or with a delay of only a few months between the peak times of the infections. Examples are influenza A and parainfluenza which have coinciding peaks, and RSV and Mycoplasma pneumoniae with a delay of about four months between the peaks [1].

Here, we explored two infectious diseases circulating during the same period, where the symptoms of only one of the diseases are severe enough to stay at home. The effect of staying at home for the disease with the severe symptoms on the final size of the other infection was studied. For two diseases with a similar basic reproduction number and a similar infectious period, staying at home for the disease with the severe symptoms can cause a small increase in the final size of the other infection in case of low basic reproduction numbers. This could be explained by a shift in the peak time of infection of the disease with the severe symptoms, resulting in a smaller number of people with less contacts at the peak time of the other infection. This effect was influenced by the mixing patterns, the timing of the two infections and the difference in basic reproduction number between the two pathogens.

Figure A.6 shows that the same effects also occur when studying a model with 86 × 86 contact matrices instead of 2 × 2 matrices, which shows that the observed effects are not an artifact of the model.

Let *R*_0,1_ and *R*_0,2_ represent the basic reproduction numbers of disease 1 and disease 2 respectively. A larger effect on the evolution of the final size of infection 2 with *p*_1_ is observed when *R*_0,1_ = *R*_0,2_ + 0.3 than when *R*_0,1_ = *R*_0,2_ − 0.3. This is because the two scenarios are not symmetric. Increasing *R*_0,1_ with 0.3 and keeping *R*_0,2_ at its value of Figure 2 causes a larger shift of the peak time of infection 1 than decreasing *R*_0,1_ with 0.3.

Second, we studied two infectious diseases for which the most severe one induces twice as many symptomatic individuals staying at home than for the other disease. Here, it was observed that no matter what the basic reproductive number is, increasing the proportion staying at home always reduces the final size of both infections, and in particular considerably reduces the number of co-infections.

Our approach has several limitations. First, variation of immunity, which can have a considerable impact on the attack rates and epidemic peaks [16], was not taken into account. Second, the study was restricted to a limited number of model variations and scenarios that were relevant to explain the effect of staying at home when having symptoms of one disease on the other, or the effect that twice as many symptomatic individuals stay at home for the most severe disease than for the other. Third, the model could be extended to more than two diseases or to other types of compartmental models such as the SEIR model (including a latent period) and the SIRS model (assuming a short period of immunity instead of life-long immunity). Fourth, we assumed that people stay at home at the onset of symptoms. In practice, people feel bad and stay home the day after. Fifth, competition between two pathogens was not taken into account. Competion could, among others, be included by assuming partial cross-immunity, or enhanced susceptibilty to one of the diseases compared to the other [4]. Sixth, the model is a non-preferential model. This means that we assume that the infection risk is the same irrespective of whether a susceptible individual is contacting a symptomatic or asymptomatic individual. Moreover, asymptomatic and symptomatic cases recover at the same rate. The model can be extended to a preferential model, like described by Santermans et al. [10] for mono-infections. Lastly, contact matrices for the 2009 A/H1N1pdm influenza from [14] were used. Using contact matrices for other strains or pathogens could influence our conclusions.

To our knowledge, this was the first study assessing the influence of changes in behavior on the joint dynamics of two infectious diseases. We can conclude that the reported effects are caused by different mixing patterns between asymptomatic and symptomatic individuals, and individuals staying at home. Furthermore, a take home message from this study is that assessing the joint dynamics of two or more infectious diseases is important to give advise on behavioral interventions. From a public health point of view, it is crucial to include age classes and differences in mixing patterns between symptomatic and asymptomatic cases in modeling studies.

## Acknowledgements

This research is part of a project that has received funding from the European Research Council (ERC) under the European Union’s Horizon 2020 research and innovation programme (grant agreement 682540 — TransMID). NH gratefully acknowledges support from the University of Antwerp scientific chair in Evidence-Based Vaccinology, financed in 2009–2017 by a gift from Pfizer and in 2016 by a gift from GSK. We gratefully acknowledge Thomas Kovac (UHasselt) for improving our R code for running the model. We thank Kim Van Kerckhove (UHasselt, Ugentec) and Eva Santermans (UHasselt, Galapagos) for their input and discussions on social contact data. We thank James Wood (UNSW Sidney, Australia) for his helpful comments and discussions that improved the manuscript.

## A System of partial differential equations

The system of partial differential equations for the model is given by:

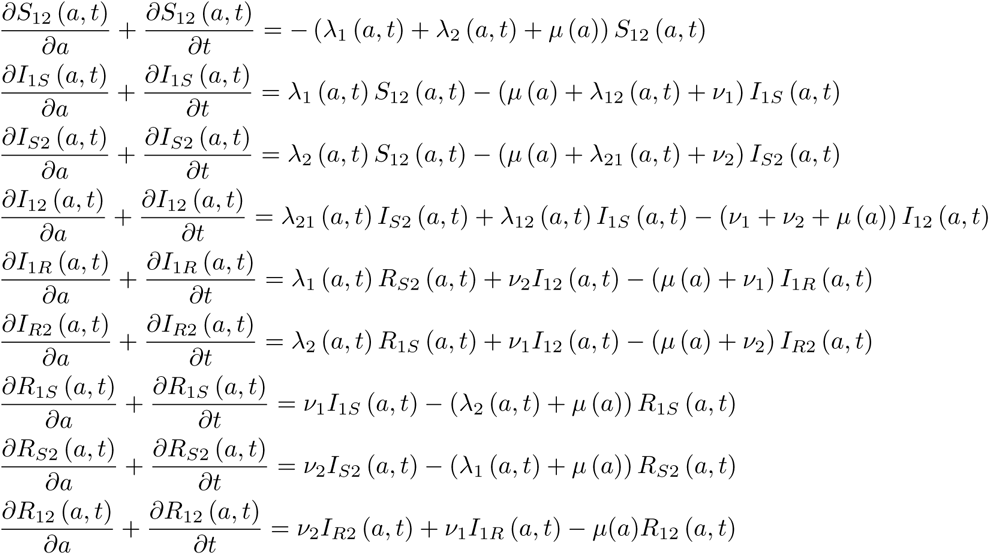

Put *I*_1_ = *I*_1_*_S_* + *I*_12_ + *I*_1_*_R_* and *I*_2_ = *I_S_*_2_ + *I*_12_ + *I_R_*_2_. For K age categories, the age-specific forces of infection are given by [6]

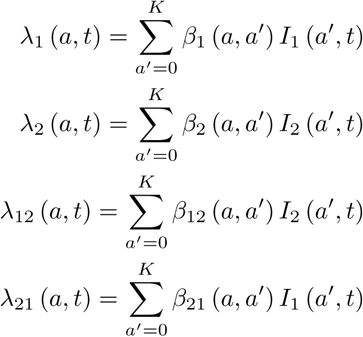

When *ϕ*_1_ (resp. *ϕ*_2_) is the proportion of symptomatic cases of infection 1 (resp. infection 2), then the age-specific transmission rates can be calculated from the contact matrices as follows [15]

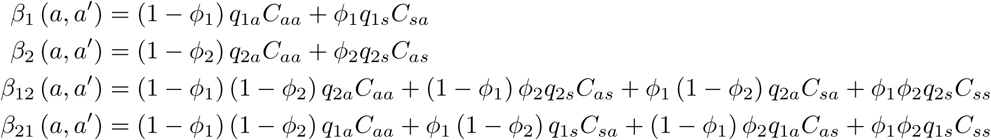

where *q*_1_*_a_*, *q*_1_*_s_*, *q*_2_*_a_* and *q*_2_*_s_* are disease-specific proportionality factors for asymptomatic infection 1, symptomatic infection 1, asymptomatic infection 2 and symptomatic infection 2. *C_aa_* is the asymptomatic mixing matrix; *C_sa_* and *C_as_* are symptomatic mixing matrices for disease 1, resp. disease 2 only; *C_ss_* is the symptomatic mixing matrix for people having symptoms for both diseases.

**Figure A.1.**
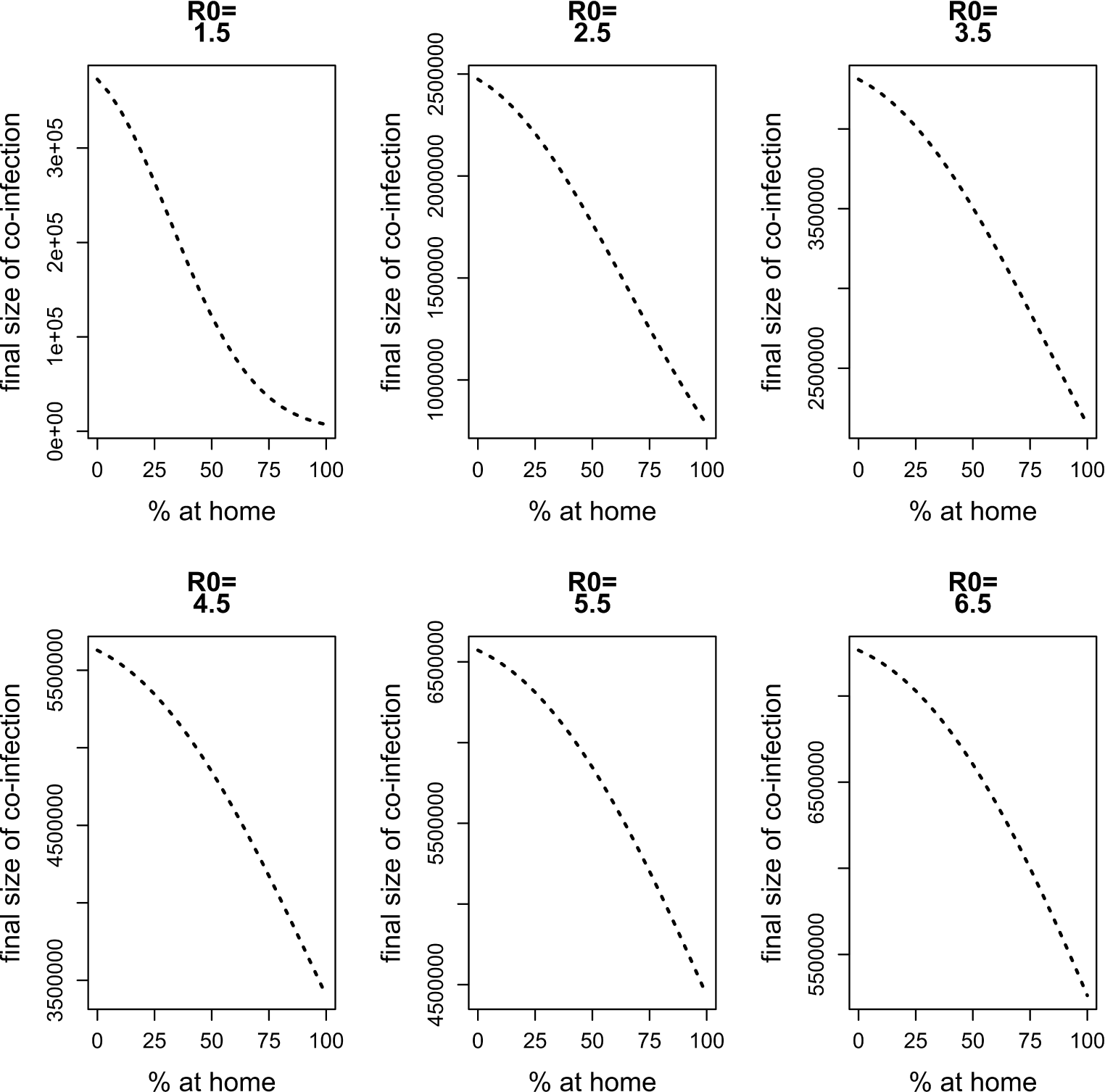
Final size of co-infection against the percentage staying at home when having symptoms of disease 1 for different values of *R*_0_.

**Figure A.2.**
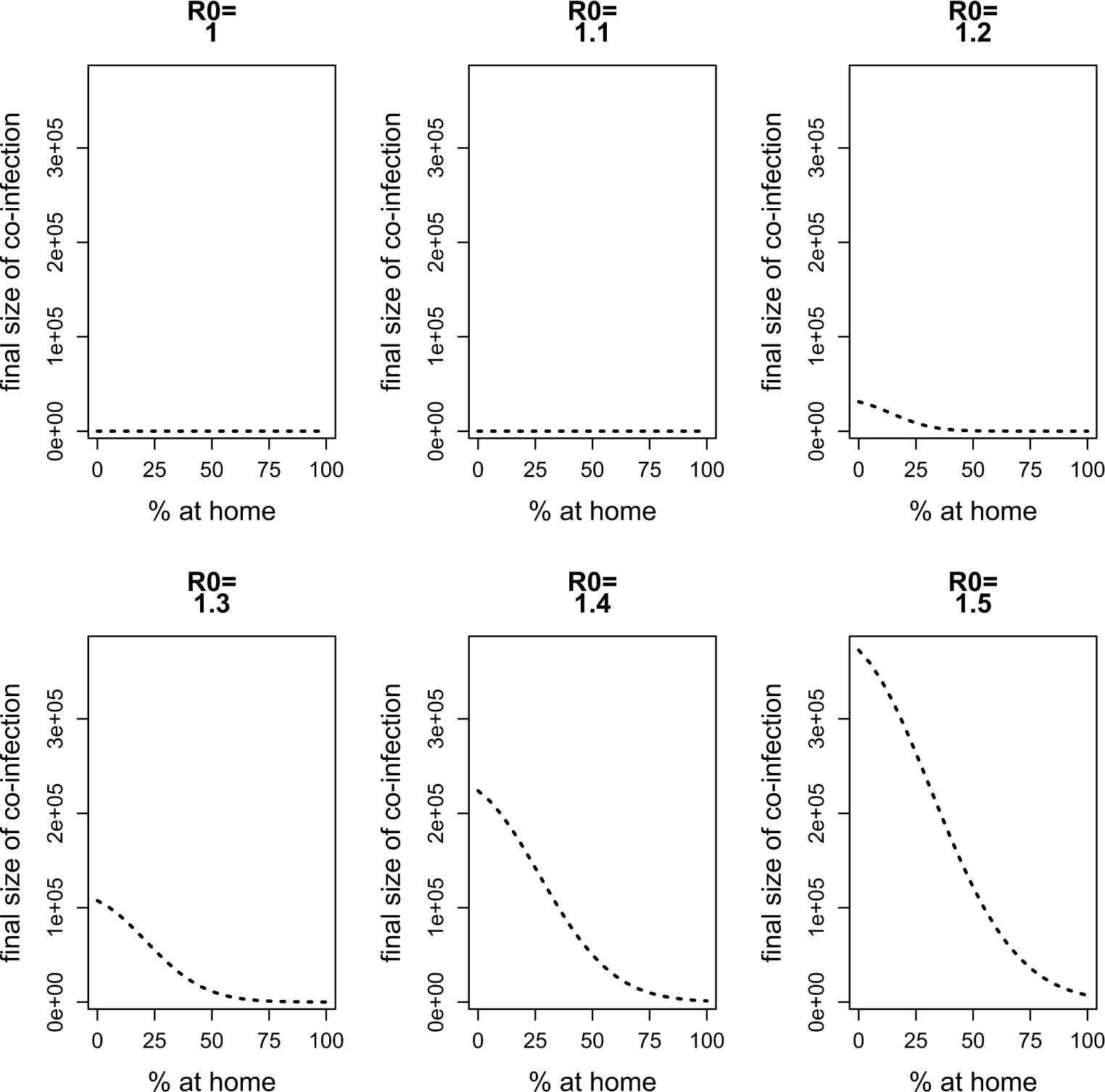
Final size of co-infection against the percentage staying at home when having symptoms of disease 1 for *R*_0_ varying between 1 and 1.5. In contrast to Figure A.1, the same scale is used on the vertical axis.

**Figure A.3.**
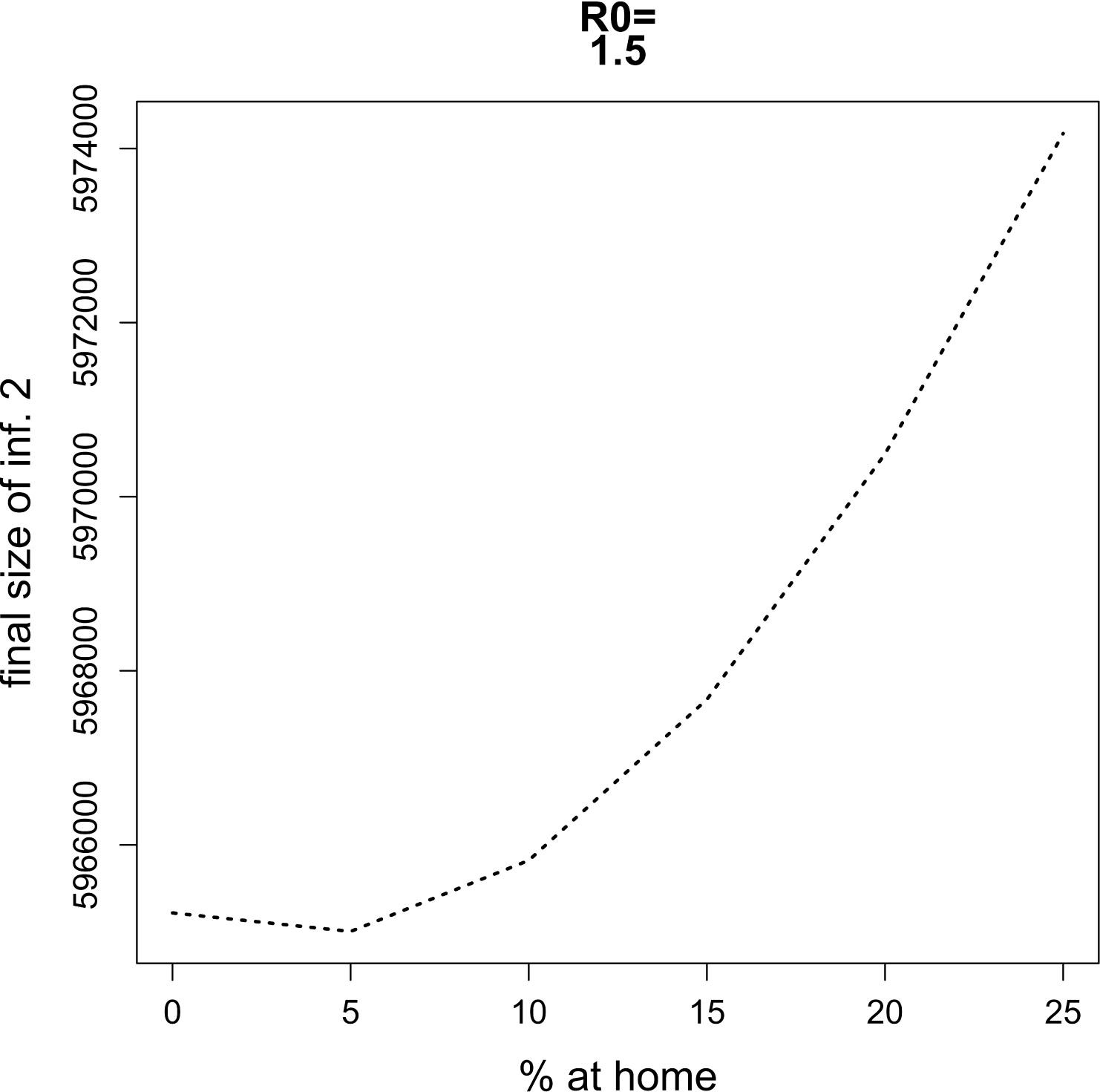
Final size of infection 2 against the percentage staying at home when having symptoms of disease 1 for *R*_0_ = 1.5 and percentages staying at home ranging from 0% up to 25% in steps of size 5%.

**Figure A.4.**
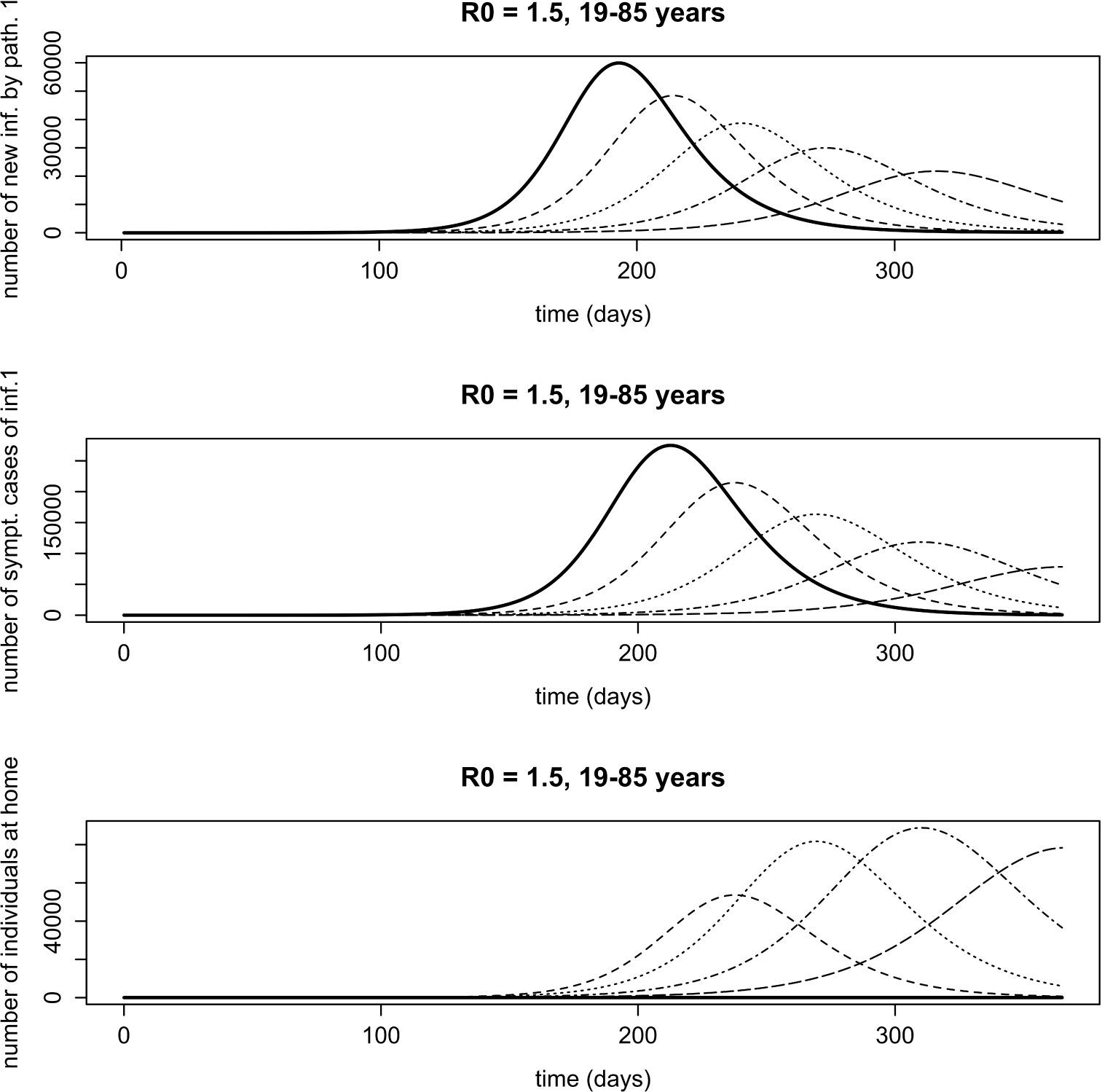
Influence of home isolation for disease 1 only, for age class 19 − 85 years in case of two infections with an infectious period of 7 days and *R*_0_ = 1.5. Solid line: 0% at home, dashed line: 25% at home, dotted line: 50% at home, dotdashed line: 75% at home, longdashed line: 100% at home. Upper panel: number of new infections with pathogen 1. Because there is no home isolation for disease 2, the number of new infections with pathogen 2 coincides with the solid line; middle panel: number of symptomatic cases of infection 1; lower panel: number of symptomatic cases of infection 1 staying at home.

**Figure A.5.**
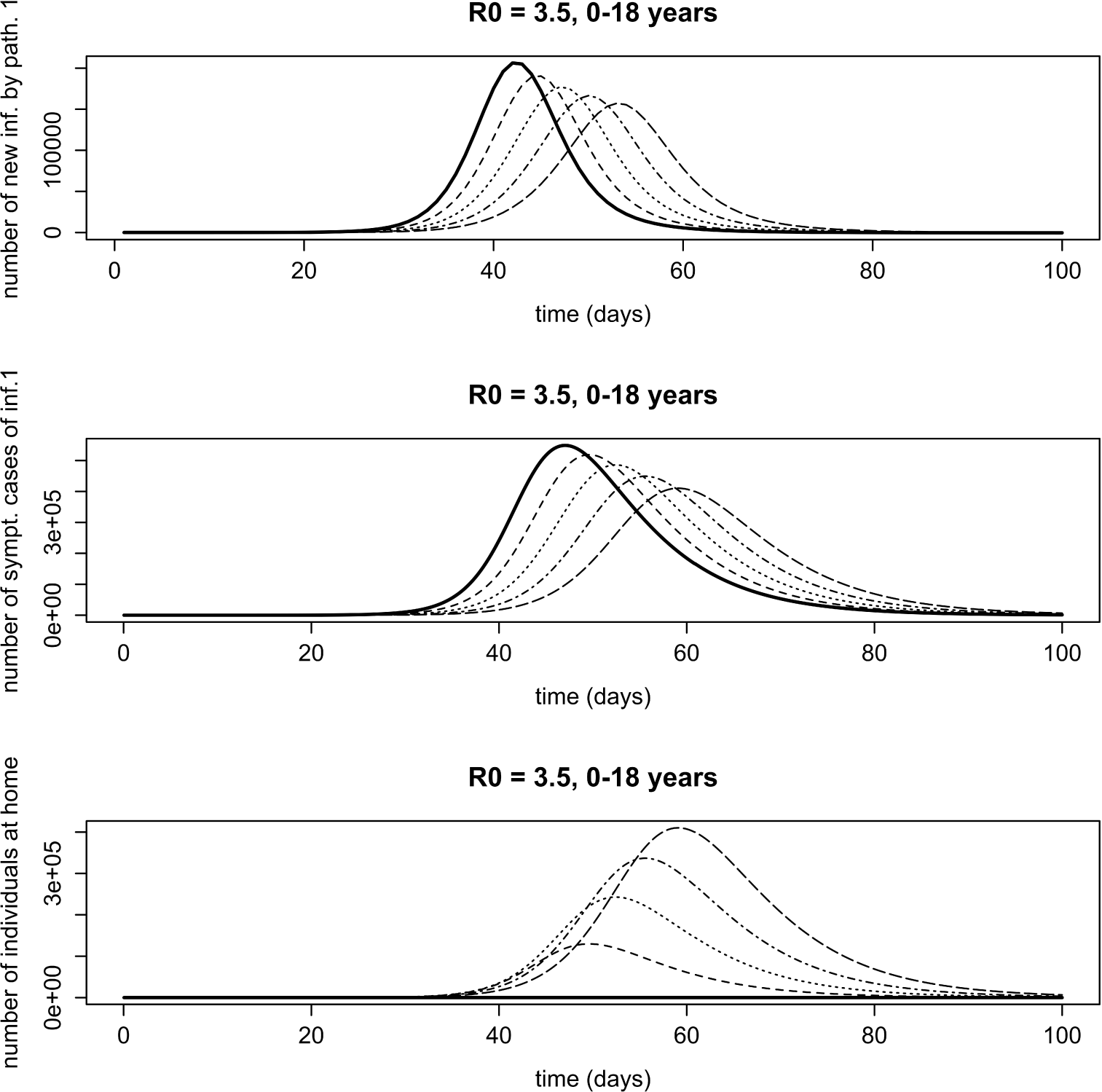
Influence of home isolation for disease 1 only, for age class 0 − 18 years in case of two infections with an infectious period of 7 days and *R*_0_ = 3.5. Solid line: 0% at home, dashed line: 25% at home, dotted line: 50% at home, dotdashed line: 75% at home, longdashed line: 100% at home. Upper panel: number of new infections with pathogen 1. Because there is no home isolation for disease 2, the number of new infections with pathogen 2 coincides with the solid line; middle panel: number of symptomatic cases of infection 1; lower panel: number of symptomatic cases of infection 1 staying at home.

**Figure A.6.**
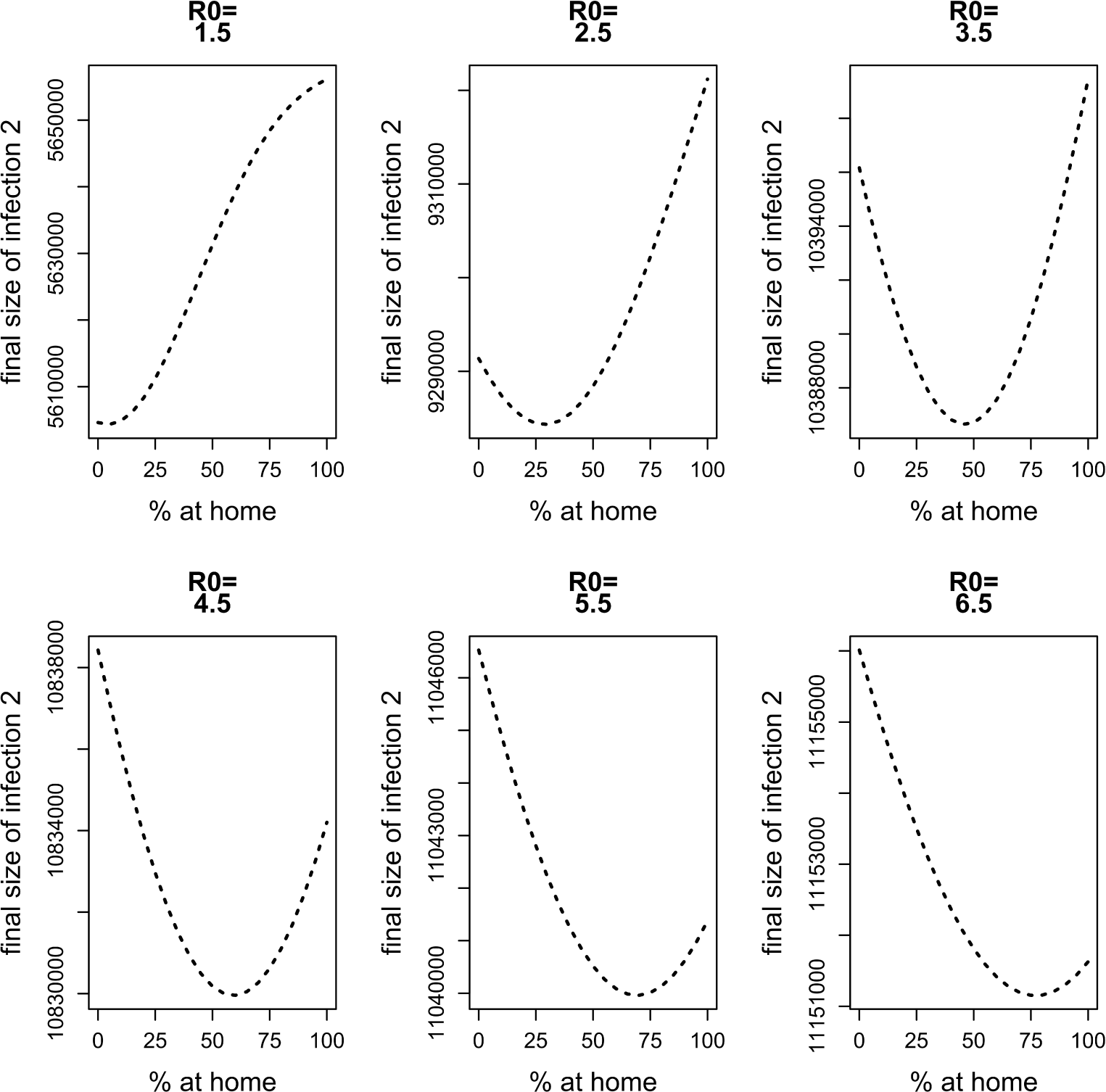
Final size of infection 2 against the percentage staying at home when having symptoms of disease 1 for different values of *R*_0_ for a model with 86 × 86 contact matrices.

**Figure A.7.**
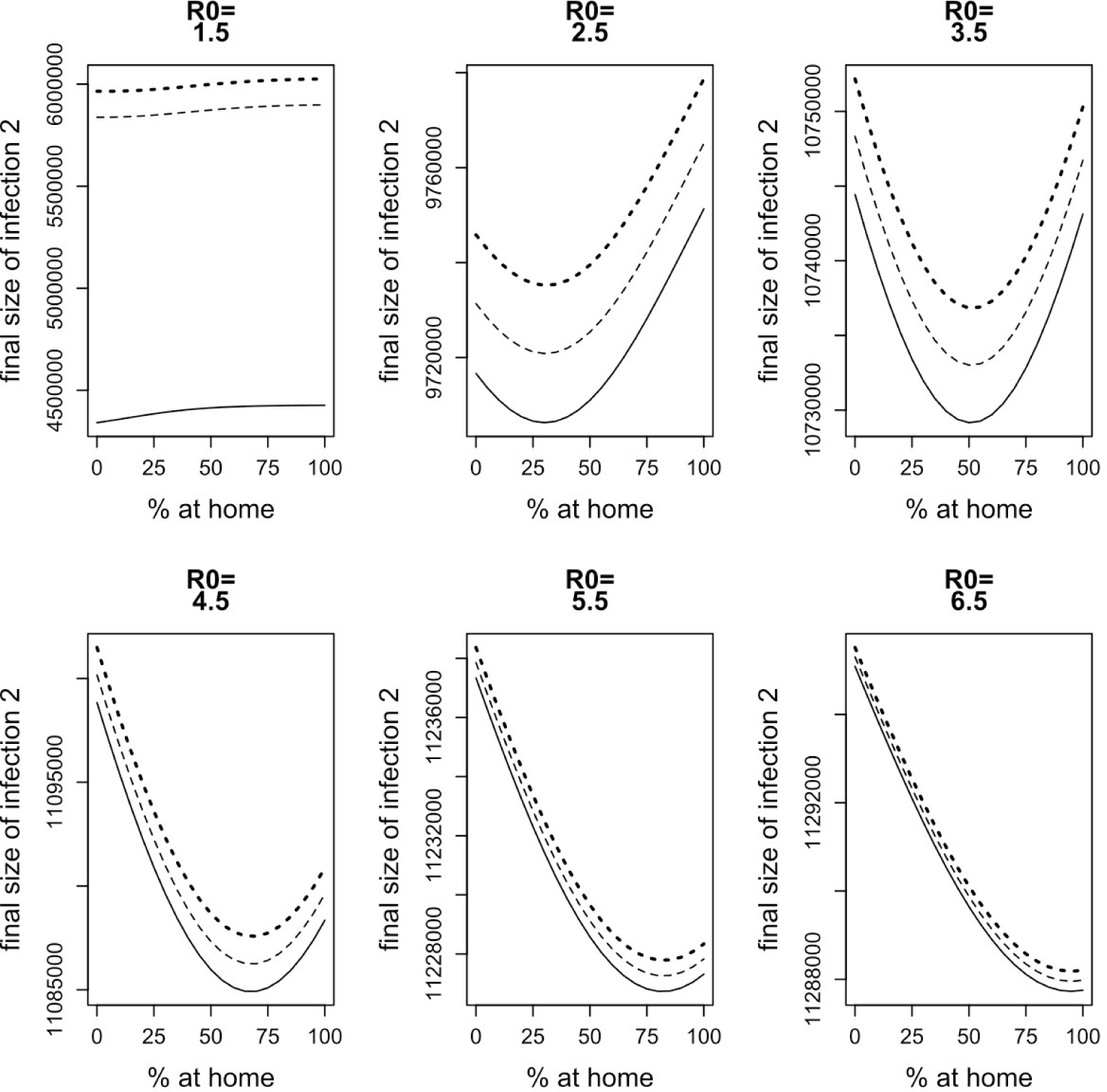
Influence of the infectious period on the behavior observed in Section 3.1. Infectious period of both infections is 7 days (dotted line); 14 days (dashed line); 21 days (solid line).

**Figure A.8.**
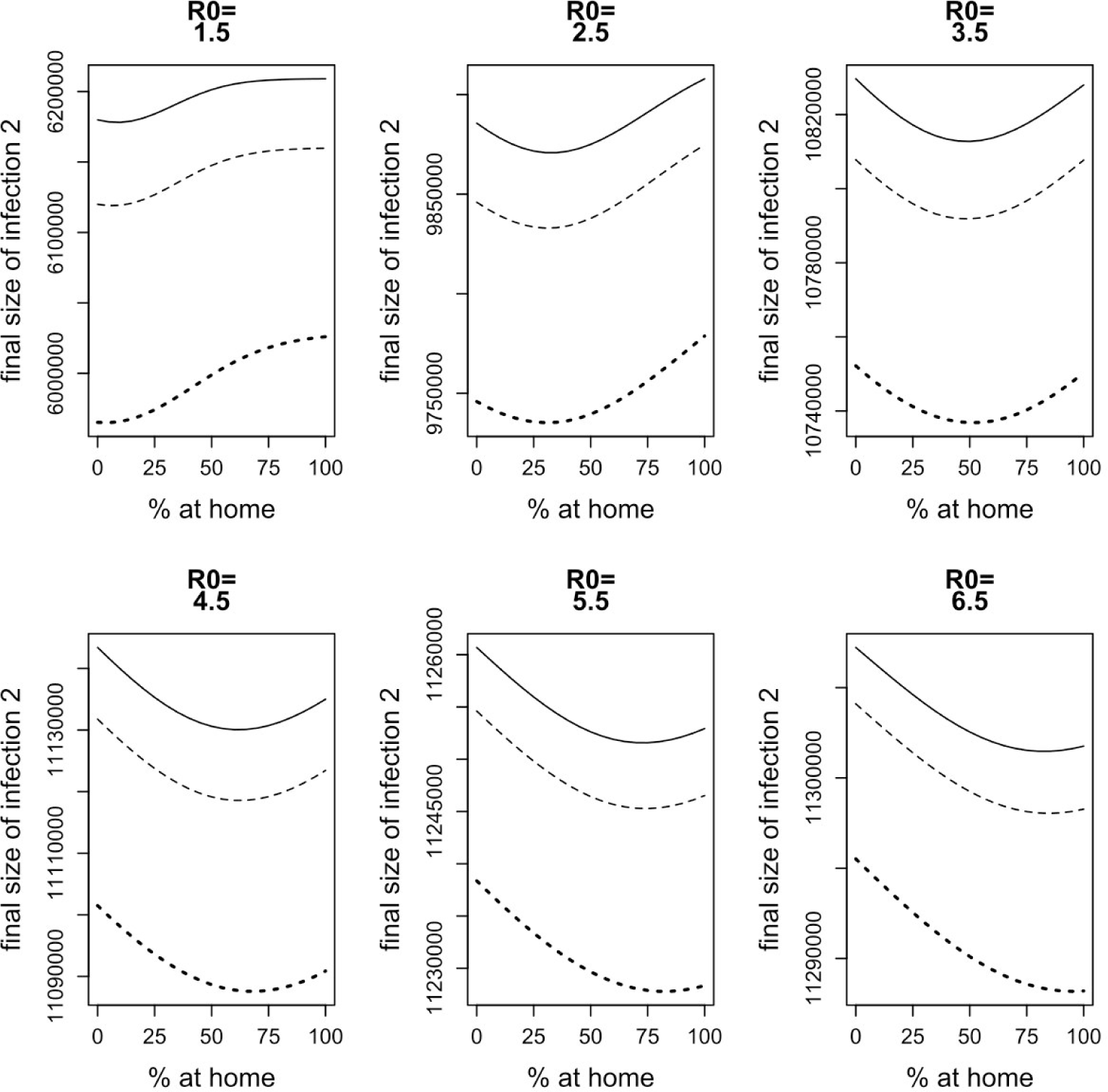
Influence of the infectiousness of symptomatic versus asymptomatic cases on the behavior observed in Section 3.1. Solid line: symptomatic cases are 9 times as infectious as asymptomatic cases; dashed line: symptomatic cases are 6 times as infectious as asymptomatic cases; dotted line: symptomatic cases are three times as infectious as asymptomatic cases.

**Figure A.9.**
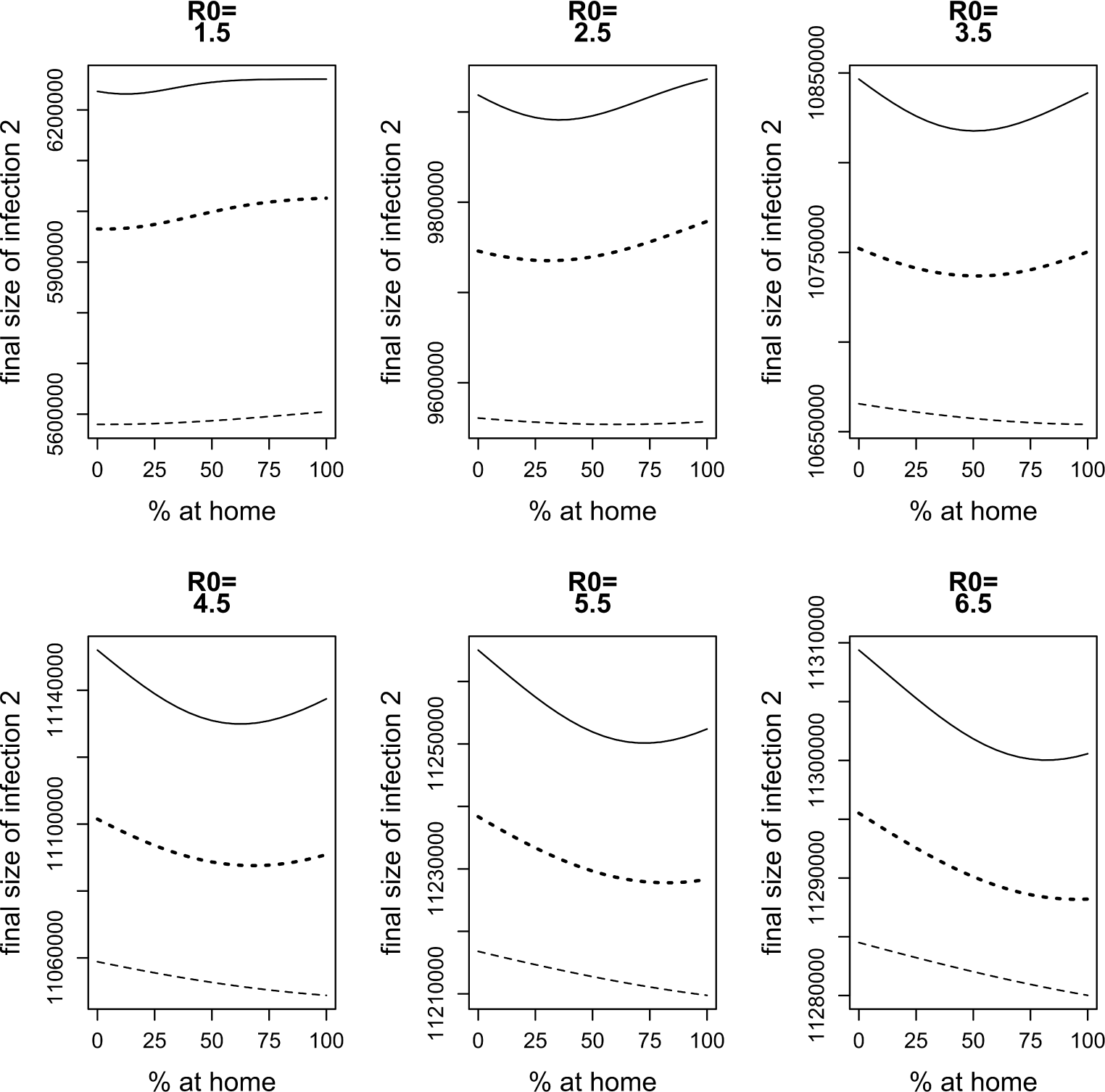
Influence of the fraction of symptomatic cases on the behavior observed in Section 3.1. Dashed line: 30% symptomatic; dotted line: 60% symptomatic; solid line: 90% symptomatic

**Figure A.10.**
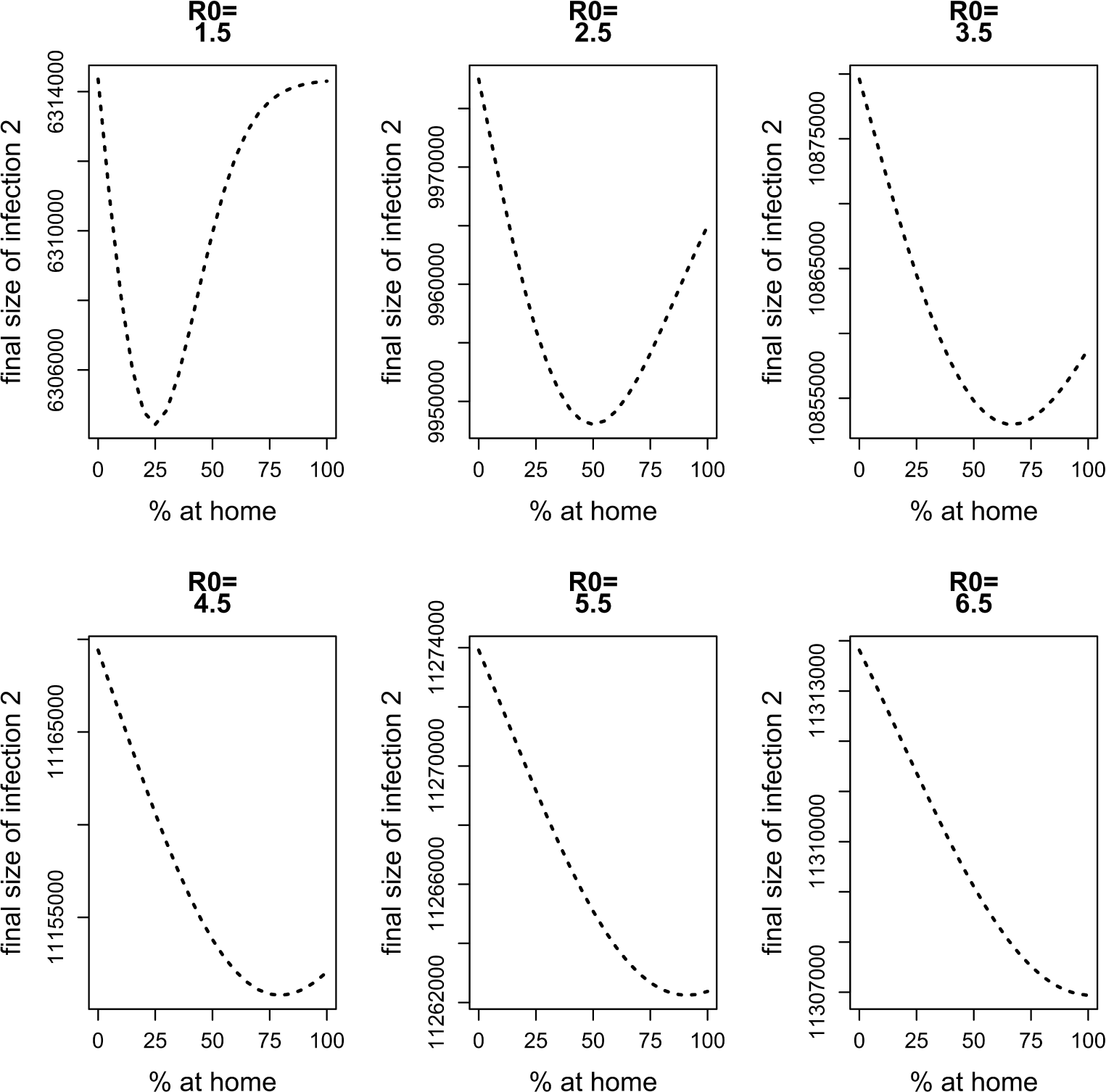
Influence of contact matrices on the behavior observed in Section 3.1. Final size of infection 2 against the percentage staying at home when having symptoms of disease 1 for different values of *R*_0_ when symptomatic individuals not staying at home and asymptomatic individuals would have the same mixing patterns.

**Figure A.11.**
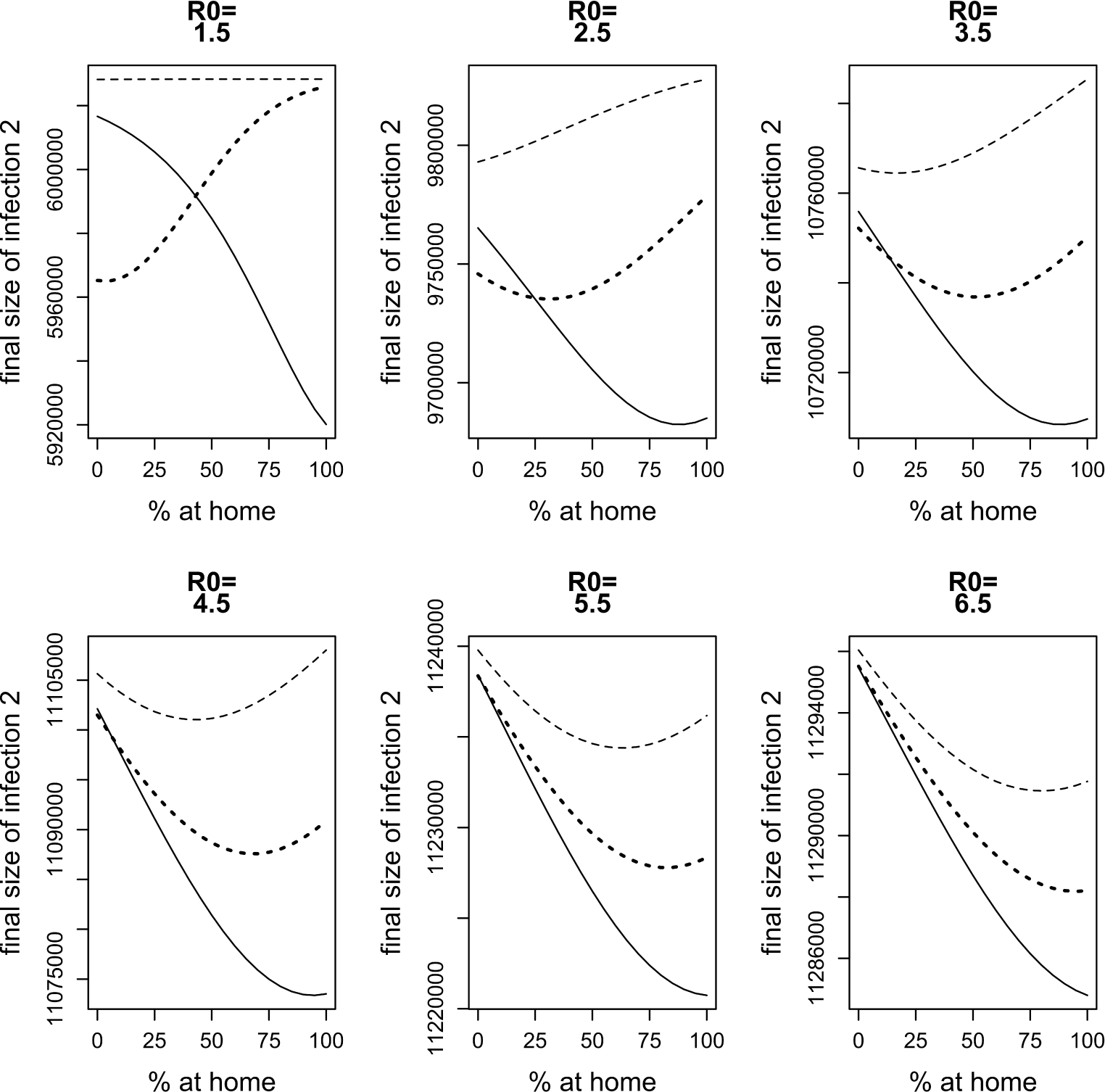
Influence of the difference in *R*_0_ between pathogen 1 and pathogen 2 on the behavior observed in section 3.1. Dotted line: *R*_0,1_ = *R*_0,2_; dashed line: *R*_0,1_ = *R*_0,2_ − 0.3; solid line: *R*_0,1_ = *R*_0,2_ + 0.3

**Figure A.12.**
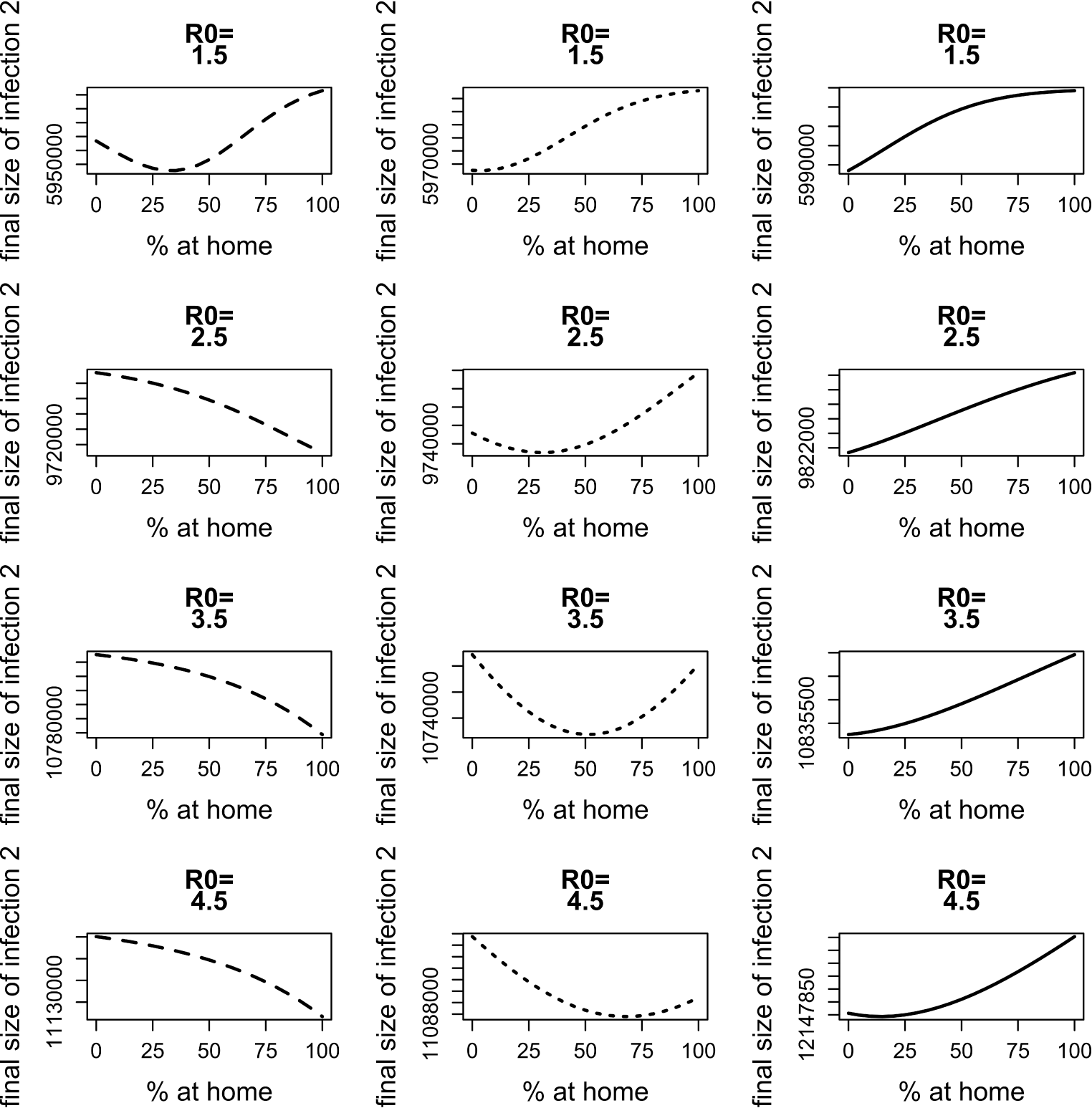
Influence of the delay between the start of the the two infections on the behavior observed in Section 3.1. Dashed line: infection 1 starts one month earlier than infection 2; dotted line: infections start at the same time; solid line: infection 1 starts one month later than infection 2.

**Figure A.13.**
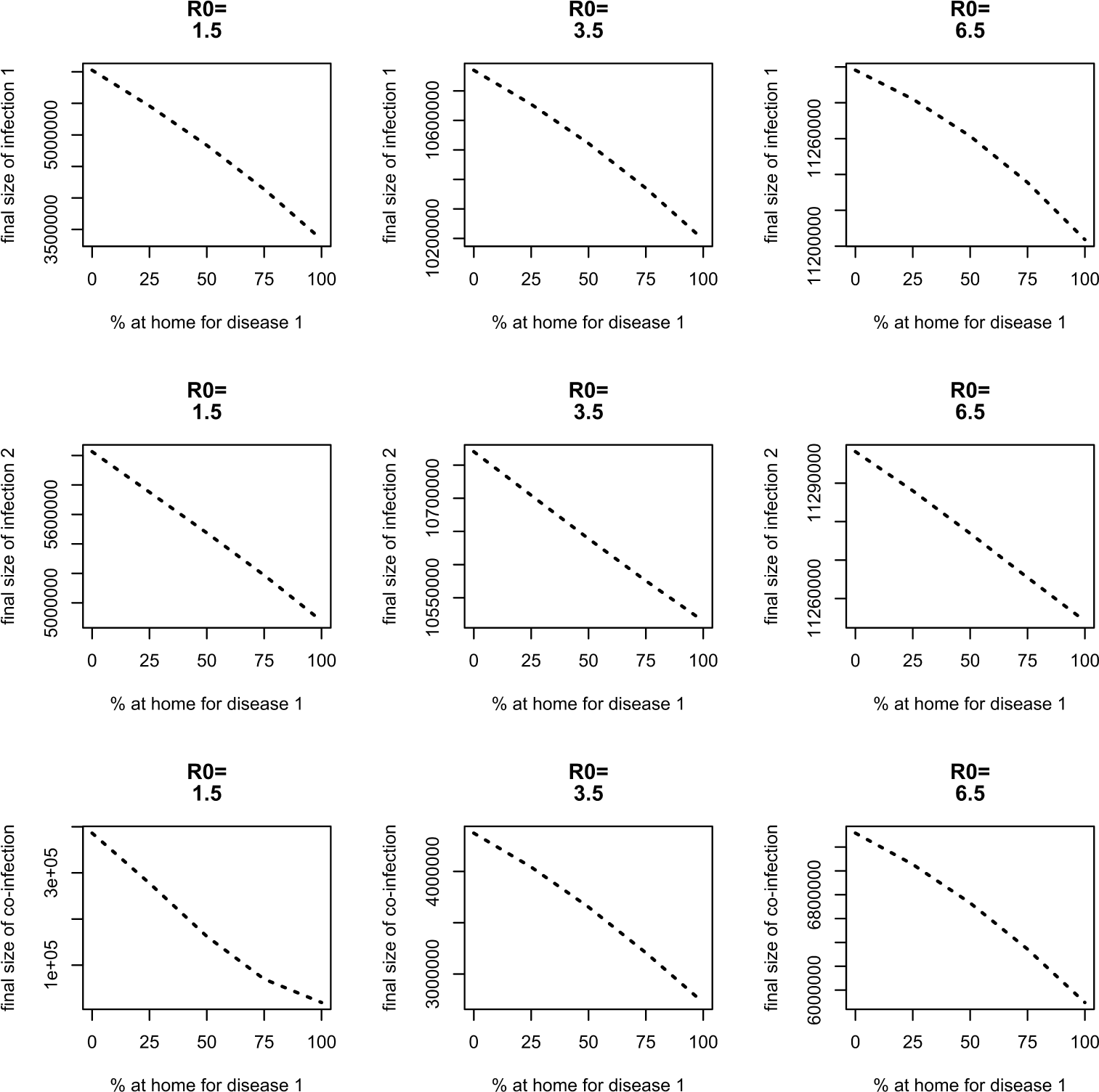
Final size of infection 1 (first row), infection 2 (second row) and co-infections (third row) against the percentage staying at home when having symptoms of disease 1 for different values of *R*_0_. Left: *R*_0_ = 1.5, middle: *R*_0_ = 3.5, right: *R*_0_ = 6.5.

